# The Structure of the Picornaviral 2C:RNA holoenzyme: Molecular Basis of RNA binding and specificity by a AAA+ protein

**DOI:** 10.64898/2026.06.07.730651

**Authors:** Richard A. Pfuetzner, Louie N. Pinpin, Maëva Duboeuf, K. Ian White, Adriana G. Tubb, Bharti Singal, David Fernandez-Martinez, William R. Arnold, Axel T. Brunger, Garvey Mckenzie, Trevor Sweeney, Yousuf A. Khan

**Affiliations:** Department of Molecular and Cellular Physiology, Stanford University, Stanford, CA, USA; Viral Gene Expression Group, The Pirbright Institute, Pirbright, UK; Department of Biochemistry and Structural Biology, Long School of Medicine, University of Texas Health San Antonio, San Antonio, TX, USA; Department of Pharmacology, Long School of Medicine, University of Texas Health San Antonio, San Antonio, TX, USA; Stanford Cryo-Electron Microscopy Center (cEMc), Stanford University, Palo Alto, CA, USA; Stanford University Mass Spectrometry, Stanford University, Palo Alto, CA, USA

**Author notes:** These authors contributed equally to this manuscript (Richard A. Pfuetzner, Louie N. Pinpin, Maëva Duboeuf).

## Abstract

Picornaviruses are one of the leading agents of animal and human infectious disease with at least 8 billion infections a year and cause a range of symptoms including respiratory failure and acute flaccid myelitis^1^. The most conserved nonstructural protein in picornaviruses is 2C^2^, a member of the AAA+ family of ATPases that binds RNA, and a broad spectrum antiviral target^3–5^. Despite its crucial role in the viral life cycle and as a clinical target, no structure of 2C bound to RNA has been structurally determined. Here we present the first structure of 2C as a hexamer bound to single stranded RNA in its central pore; a novel AAA+ protein:substrate interaction. Using the 2C:RNA holoenzyme complex structure, we characterize the mode that this AAA+ protein employs to specifically bind single stranded RNA, and demonstrate that mutations to key residues inhibit both RNA binding and viral replication in *Apthovirus* and *Enterovirus* systems, and show that the core residues responsible for binding are broadly conserved in viruses beyond *Picornaviridae*. Finally, we reveal that the 2C:RNA holoenzyme complex is conformationally more similar to a protein translocase adapted to bind RNA rather than other viral DNA binding SF3 helicases, underscoring how the AAA+ core module can be adapted for a variety of biochemical substrates.

## Main

The *Picornaviridae* family, or picornaviruses, are non-enveloped, positive-sense, single-stranded RNA viruses that cause a wide range of highly infectious diseases in humans and animals^6^. In humans, picornaviruses are responsible for at least 8 billion infections a year^1^ that can lead to a swath of diseases ranging from upper respiratory failure to paralysis^7^. The economic burden of viral respiratory tract infections alone in the United States of America caused by picornaviruses is estimated to be approximately $40 billion^8^. In animals, foot-and-mouth disease virus (FMDV) was the first animal virus discovered^9^ and foot-and-mouth disease in livestock results in approximately $21 billion loss globally^10^, while avian picornaviruses also lead to disease in poultry^11^. Lastly, recent outbreaks of different enteroviruses have led to acute flaccid myelitis (EV-D68^12^) or paralysis and death (EV-A71^13^) in children globally. Despite the immense impact to human and animal health with a death toll that is poorly quantified globally and across all different types of picornaviruses, there are few antiviral therapies approved for clinical use against non-polio picornaviruses^14^.

2C is the most conserved nonstructural protein in picornaviruses^2^ and is therefore an attractive target for the design of broad spectrum antivirals^3–5^. Critically, 2C is required for viral replication yet its many potential functions, including putatively packaging the viral RNA into the capsid^15–17^, and mechanism of action remain poorly understood^18^. The presence of a conserved AAA+ (<u>A</u>TPases <u>a</u>ssociated with diverse cellular <u>a</u>ctivities^19^) domain with canonical Walker A, Walker B, Motif C, and arginine finger motifs to coordinate ATP binding and hydrolysis has led to its classification as a clade 4 AAA+ SF3 helicase^20^, along with the DNA helicase SV40 T-antigen^21^, though previous work has shown that 2C binds to single stranded RNA (ssRNA)^18,22,23^. This indicates that 2C, akin to other SF3 helicases and AAA+ complexes such as the proteasome^24^ and protein translocases^25–27^, likely forms a hexameric structure in its active form to associate with its RNA substrate. Yet, no studies have been able to capture the structure of this active hexameric holoenzyme complex, which would provide insight into the first putative AAA+ complex to bind RNA, help understand key mechanistic principles by which this complex functions, and serve as the basis for the development of broad viral inhibitors.

In this study, we resolve the FMDV 2C hexameric complex bound to ssRNA by electron cryogenic microscopy (cryoEM) in which we capture the protomers in several different nucleotide states and with a large split in the ring. We were able to resolve a ssRNA bound directly in its pore which, according to our knowledge, is the first observed structure of any AAA+ protein bound to RNA. We found that the 2C:RNA holoenzyme complex employs a varied network of interactions to bind to RNA all through its central pore and that these residues in 2C are conserved in other picornaviruses as well other RNA viruses beyond the *Picornaviridae* family that contain a 2C-like AAA+ ortholog in their genome. Mutations to these residues reduced binding of 2C to ssRNA *in-vitro* and inhibited replication in FMDV and coxsackie virus B3 (CVB3) systems. Lastly, we show that 2C phylogenetically clusters between other SF3 helicases and AAA+ protein translocases and contains key AAA+ structural features that are more characteristic of a protein translocase than an SF3 helicase underscoring how the core AAA+ module can be manipulated for different substrates and functions.

### The 2C:RNA holoenzyme complex

Picornaviral structural and nonstructural proteins are released from a polyprotein by viral protease cleavage^28^ (**Figure 1a**). 2C possesses an N-terminal amphipathic helix responsible for membrane association^29^, a helical bundle domain^30^ and a C-terminal AAA+ domain^31^. For structural studies, we used a construct with containing FMDV 2C with the N-terminal amphipathic helix truncated and an N207A mutation in Motif C to slow ATP hydrolysis (**Extended Figure 1, Materials and Methods**)^23^. This mutant bound ssRNA (**Figure 1b**) with no comparable difference to the WT (**Extended Figure 2a-b**). Mixing Δ33 2C N207A with ssRNA in the prescence of ATP and Mg^2+^ yielded a large molecular weight species that appeared to form a complex in negative stain cryoEM (**Extended Figure 2c-e**). We then obtained 260,225 high-quality particles (**Extended Figure 2f, Extended Figure 3**) that yielded a final 3.05Å map.

**Figure 1:**
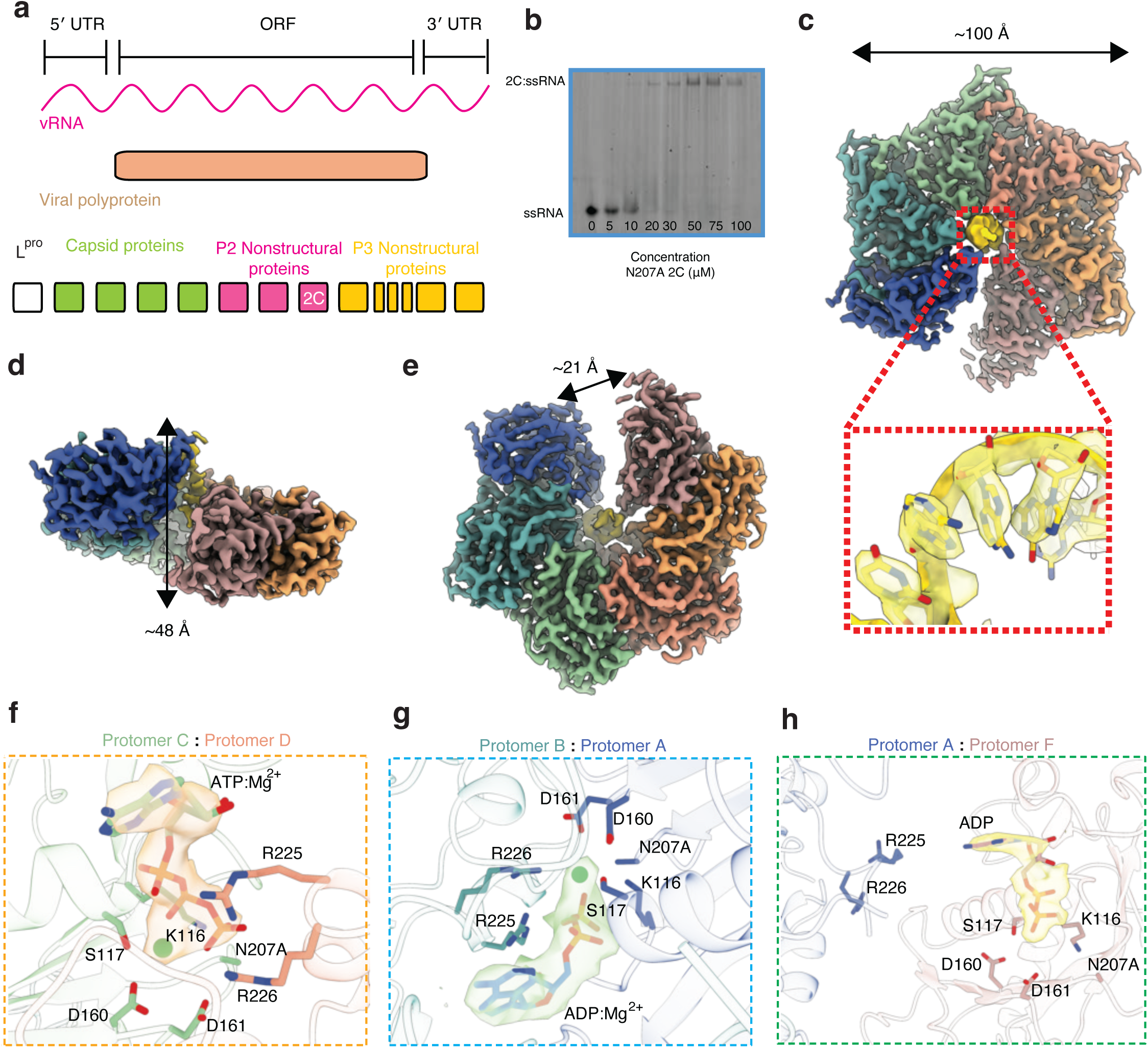
Architecture of the 2C:RNA holoenzyme complex: **a.** Schematic of the viral RNA to the final cleaved viral products, including 2C, for FMDV. **b.** EMSA of Δ33 2C N207A with ssRNA at varying concentrations of protien. **c.** Top view of 2C cryoEM map colored by protomer with inlet of density (yellow) surrounding ssRNA substrate. **d.** Side-view of split ring of 2C. **e.** Bottom view of 2C cryoEM map with split ring. **f.** Protomer interface between Protomer C (green) and Protomer D (orange) with density highlighting the ATP and Mg^2+^. **g.** Protomer interface between Protomer B (dark green) and Protomer A (dark blue) with density highlighting the ADP and Mg^2+^. **h.** Protomer interface between Protomer A (dark blue) and Protomer F (mauve) with density highlighting the ADP.

The 2C:RNA holoenzyme complex consists of six identical 2C protomers to form a hexameric ring complex with a diameter of 100Å (**Figure 1c**). Like other AAA+ complexes, we found the protomers to form a spiral staircase around the axis defining its central pore^19,32^ (**Figure 1d**) and a large split between its top (protomer A) and bottom (protomer F) protomers reminiscent of AAA+ protein translocases^19,25,26,33^ (**Figure 1e**). This is in stark contrast to other SF3 helicases that typically form a tight, closed planar ring around their DNA substrate^21,34^. Most notably, we were able to find density of ssRNA bound to the 2C central pore with an approximate local resolution between 2.5 Å and 3.0 Å at or proximal to the RNA, allowing us to model the first instance of a AAA+ protein bound directly to RNA as its substrate (**Extended Figure 4a-b**). We also found density corresponding to the “collar” or helical domain in our cryoEM map (**Extended Figure 4c**) but this region is likely flexible as further refinement methods (particle subtraction, local refinement, etc) was unable to resolve it to a resolution that would allow for confident docking or model building.

### Interprotomer interface

As we assembled the 2C:RNA holoenzyme complex in hydrolyzing conditions (i.e. in the presence of ATP and Mg^2+^), we in principle imaged the system in a state consistent with substrate processing^25,26,35,36^. We found that the individual protomers were all bound to nucleotide but in varying states depending on the protomer (**Extended Figure 4d**). Protomer A and protomer F, both at the split, have ADP bound while protomers B-E have ATP bound. This pattern of four ATP and two ADP state (at the seam) is seen in other AAA+ protein translocases such as the proteasome^24^ or PEX1/6^37^ whereas the SV40 T-antigen SF3 helicase has been shown to transition between five ADP and one ATP to four ADP and two ATP during DNA unwinding^21^.

Close inspection of interprotomer interactions around the ATP-binding site reveals the Walker A motif, K116 and S117, and Walker B motif, D160 and D161 (**Figure 1f**) supported by the arginine finger, R225 and R226, from the neighboring protomer binding. Motif C, N207 (Ala in this structure), is also positioned proximal to the γ-phosphate. The nucleotide interface between Protomers B and A, positioning an ADP between them, is highly similar to the other interfaces positioning an ATP between them (**Figure 1g**). At the seam or split in the 2C:RNA holoenzyme complex, between protomers F and A, we found density for an ADP molecule that was only bound to Protomer F with protomer A’s arginine finger too far to bind to the nucleotide or its binding pocket (**Figure 1h**).

While the nucleotide binding pocket interactions contribute to holding AAA+ complexes together^19,32,33^, we identified a hydrophobic pocket between the protomers, closer to the substrate pore than the nucleotide binding pocket on the top face of the complex (**Extended Figure 4e**). In this pocket, we found a series of aromatic and hydrophobic residues lining the interface between protomers to form a greasy pocket with one protomer donating W139, Y140, P142, P143 and P145 and the other providing Y173, F183, I184, P185, and P186 to zipper the interface. Despite the protomers existing in different nucleotide states, the RMSD between protomers when aligned was less than 1Å (**Extended Figure 4f**).

### Mechanism of RNA binding and specificity

Inside the hexameric complex’s central pore, we found density corresponding to the ssRNA that was mixed with Δ33 2C N207A (**Figure 1c**, **Figure 2a, Extended Figure 5a-c**). We determined the sequence of the ssRNA within the pore, as the resolution of our reconstruction enables us to distinguish purines from pyrimidines, using a unique sequence within the ssRNA oligonucleotide we used for assembly of the 2C:RNA holoenzyme complex (**Extended Figure 6a-b**). Within the central axis of the pore, each protomer interacts with the substrate ssRNA in an extended fashion (**Figure 2a, Extended Figure 6a-g**); each protomer spans ∼8 nucleotides of ssRNA (**Figure 2b**). At the top of the protomer L190 packs against the hydrophobic π-interface of nucleobases and K193 hydrogen bonds with the phosphate backbone. As with other AAA+ complexes, we find an aromatic side chain, H147, that intercalates against the backbone of the substrate (**Figure 2b-c**) in a staircase-like arrangement. K169 and D146 are also in proximity to the RNA backbone for all protomers (**Figure 2a-b**). K169 hydrogen bonds directly to the nucleobase at the Watson-Crick interface (**Extended Figure 6e**) but D146, as a hydrogen bond acceptor also facing the Watson-Crick interface, does not appear to directly bind to the RNA but may serve a role in stabilizing the interactions (**Extended Figure 6c**). The density of the RNA is more diffuse near the bottom of the pore but clearly present, demonstrating that RNA runs through the entire length of the pore (**Extended Figure 5a-b**). We find at the base of the pore, or the ‘bottom’ of the complex, that R215 is proximal to this more diffuse RNA density (**Extended Figure 5c**).

**Figure 2:**
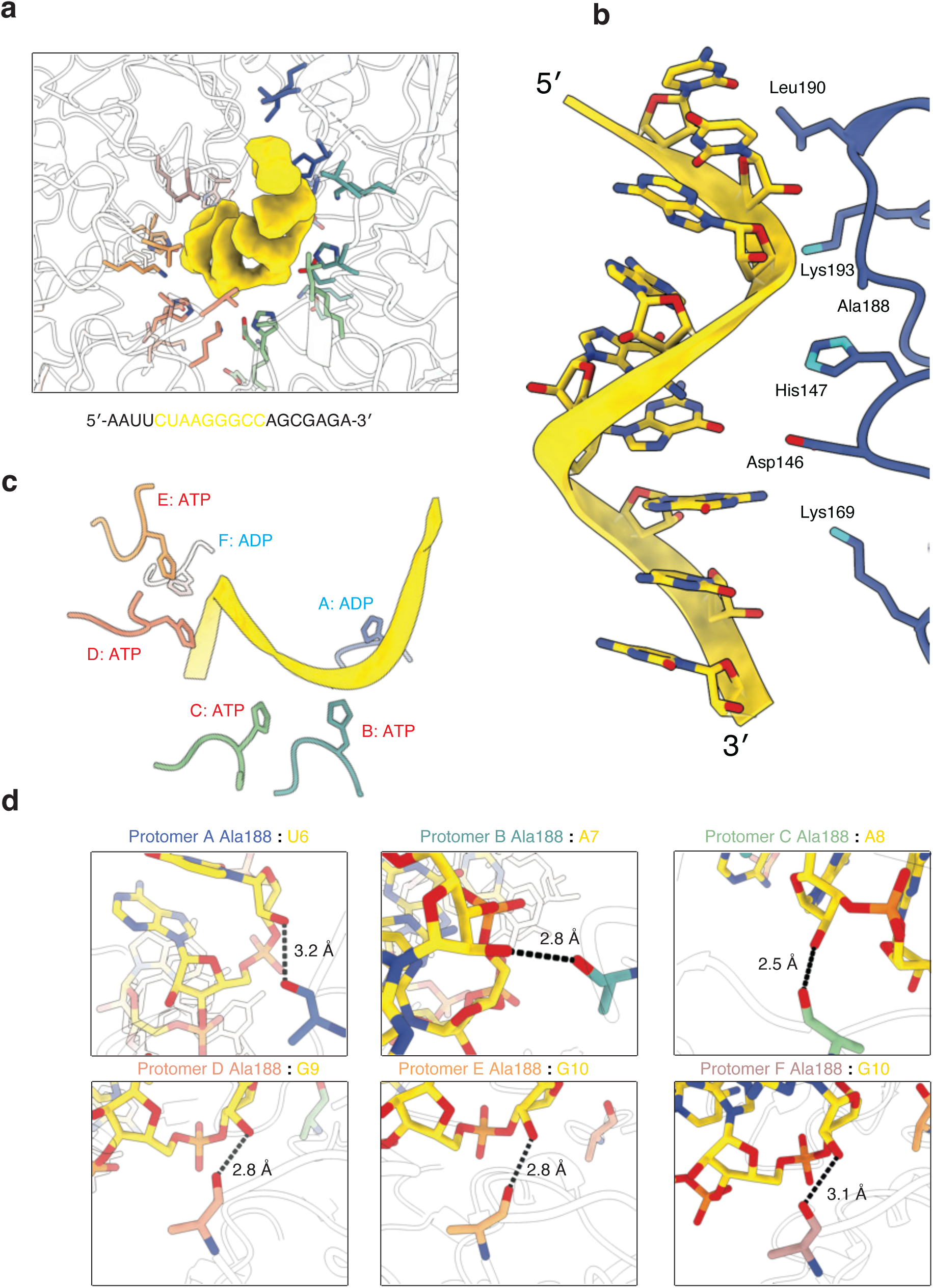
2C forms an extended ladder of interactions with substrate ssRNA: **a.** Top view of pore with residues within ∼4Å of the ssRNA along all protomers. The RNA density is shown in gold. **b.** Schematic of a single protomer’s residues within ∼4Å of the ssRNA model. **c.** Intercalating interactions of H147 along the ssRNA. **d.** Substrate specificity conferred by carbonyl’s of A188 in protomers A-F.

Interestingly, Ala188 is positioned within the pore at every protomer. While Ala188 is uncharged and has a relatively inert side chain, we find that it is not the side chain that is positioned towards the pore but rather its carbonyl oxygen (**Figure 2d**). In every protomer, the carbonyl oxygen is positioned such that it can form a moderate to strong hydrogen bond as a hydrogen bond acceptor for the 2′-OH (**Figure 2d**). Not only does this serve to bind RNA but it is the only interaction with the substrate that is unique to RNA as opposed to a hypothetical ssDNA substrate, providing a likely mechanism for 2C’s preference for RNA over DNA binding^38^.

To determine the contribution of these residues to RNA binding, we generated point mutants at the aforementioned positions and then performed protein:RNA electrophoretic mobility shift assays (EMSAs) (**Figure 3a-g**). All RNA-binding mutants expressed and purified comparably to the WT and N207A mutant. D146A had comparable binding to WT and N207A 2C, consistent with a lack of direct interaction with the ssRNA (**Figure 3a, Extended Figure 2a**). Surprisingly, replacing H147, that can both hydrogen bond or serve as a steric intercalator with the substrate, with Ala (H147A) also resulted in WT-like binding affinity (**Figure 3b, Extended Figure 2b**). This result suggests that H147 is not involved with RNA binding per se but may aid with its translocation, as is the case for other AAA+ protein translocases that still bind their substrate when their conserved pore loop aromatic amino acid is mutated but has reduced biochemical activity^39–41^. All other mutants showed decreased binding, with the largest decreases coming from A188S, L190D, and K193A (**Figure 3c-g**).

**Figure 3:**
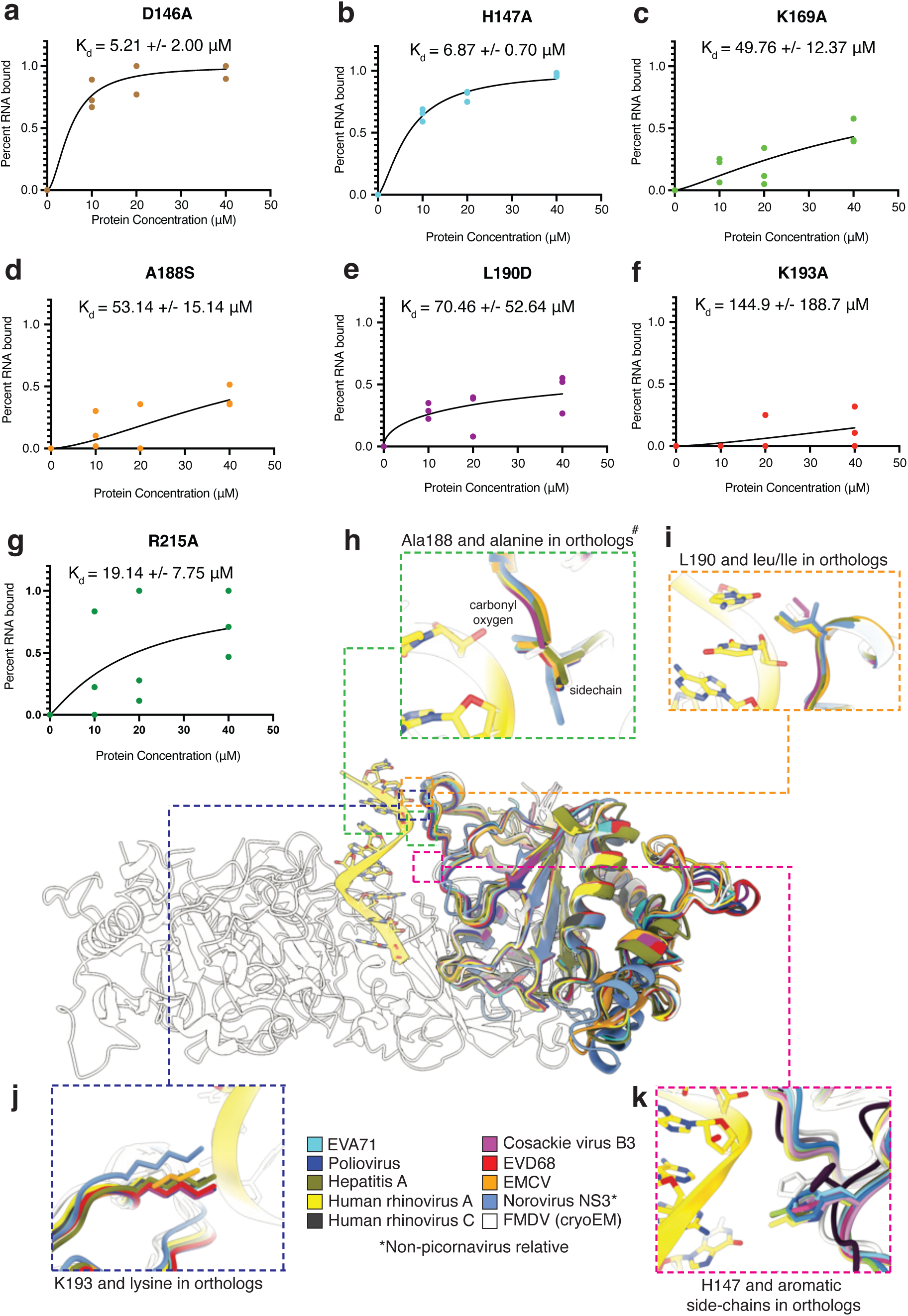
Mutations to 2C’s binding modality to ssRNA highlights evolutionarily conserved residues: EMSA binding data of ssRNA to different mutants of 2C (**a-g**) where n=3 and percent RNA bound was determined by gel densitometry. This consists of **a.** D146A, **b.** H147A, **c.** K169A, **d.** A188S, **e.** L190D, **f.** K193A, and **g.** R215A. Alignment of colabfold monomer predictions 2C and 2C ortholog (NS3) to 2C:RNA holoeznyme model. **h.** Zoom in on Ala188 and orthologs. **i.** Zoom in on L190 and orthologs. **j.** Zoom in on K193 and orthologs. **k.** Zoom in on H147 and orthologs. # Indicates a non-alanine ortholog but with the carbonyl positioning still structurally conserved.

### A conserved binding modality beyond Picornaviridae

To determine how conserved this ssRNA binding modality is, we generated colabfold^42^ predictions of the 2C monomer from other picornaviruses such as human rhinovirus A, the most frequent cause of the common cold^43^ and EVD68, an enterovirus linked to several recent outbreaks causing neurological disorders such as acute flaccid myelitis^44^. We also included the NS3 protein from Norovirus, a member of the Caliciviridae family of viruses and the most common cause of viral gastroenteritis^45^; NS3 is also a viral SF3 AAA+ ATPase related to 2C^30^.

We aligned these predictions to our 2C:RNA holoenzyme complex (**Figure 3h-j**). A188’s positioning of its carbonyl was found to be broadly conserved in our selected 2C variants and relatives (**Figure 3h**); in the case where the aligned amino acid was not an alanine, the carbonyl oxygen was still well aligned to hydrogen bond with the 2′-OH and its side chain was oriented away from the RNA-binding pore. L190 was also found to be structurally conserved, with orthologs only comprising leucine or isoleucine residues in a similar structural pose (**Figure 3i**). K193 was found to be well conserved with orthologs all containing a Lys at the same structural position (**Figure 3j**). Finally, although H147 showed no decrease in RNA binding, it was found that aromatic residues were structurally conserved at the same position among orthologs(**Figure 3k**).

Given this structural conservation of the core binders and interactors to substrate, we performed a broad structural phylogenetic search for 2C orthologs against the entire order of *Picornavirales* using the model we built into our cryoEM map. We found 497 significant hits (**Extended Figure 7**) for orthologs in the *Picornaviridae, Dicistroviridiae, Calciviridae, Iflaviridae, Secoviridae, Noraviridae, Solinviviridae, Marnaviridae*, and *Polycipiviriade* families and generated an alignment (**Extended Figure 8a**). The regions that showed the best alignment and conservation were the Walker A and B motifs, followed by the Motif C and arginine fingers regions. The C-terminal bundle was the least conserved region. Interestingly, we found that a bulky aromatic residue was conserved at the same or adjacent positions as H147 (**Extended Figure 8b**). K169 and R215 were not well conserved but A188, L190 and K193 were well conserved at their respective alignment position or at adjacent positions (**Extended Figure 8c-f**).

Given the structural and phylogenetic conservation of the binding triad (A188, L190, and K193) as well as H147, we wanted to further assess these mutants’ effect on the function of 2C in viral replication. Because 2C is a critical protein in the replication of viral RNA, we assessed replication in both a FMDV (*Apothovirus*) replicon system and an infectious CVB3 (*Enterovirus*) viral system (**Figure 4, Materials and Methods**). For both, viral replication is assessed by quantification of GFP, encoded in the respective polyprotein open reading frames. Consistent with our RNA-binding and conservation data, FMDV replicon 2C D146A and its CVB3 ortholog D162A showed no difference in replication compared to WT (**Figure 4c,e**). However, in contrast with our RNA-binding data, but consistent with conservation, we found that FMDV 2C H147A and its CVB3 2C ortholog H162A fully inhibited viral replication in both the FMDV replicon and CVB3 viral systems, respectively, demonstrating that H147 is critical in 2C’s function but not to its direct binding of RNA. All other mutants (in the FMDV replicon) system showed no replication, comparable to a mock transfection (**Figure 4d**). In the fully replicating enteroviral CVB3 system, there was no equivalent for R215 as additionally evidenced by its lack of conservation (**Extended Figure 8f**). Mutation of the conserved core binding residues all inhibited CVB3 replication (**Figure 4f**). Interestingly, the only mutant that showed a difference in effect between systems was 2C K185A in the CVB3 system (K169A ortholog in FMDV). We saw no appreciable difference between the mutant and WT CVB3 virus even though the equivalent substitution inhibited replication in FMDV. Perhaps, as suggested by the phylogenetic data (**Extended Figure 8c**), that this residue is not a core RNA binding residue for 2C across different systems. In summary, the three mutations that showed the largest reduction in ssRNA binding plus H147A all inhibited 2C replication (**Figure 4c-f, Extended Figure 9**).

**Figure 4:**
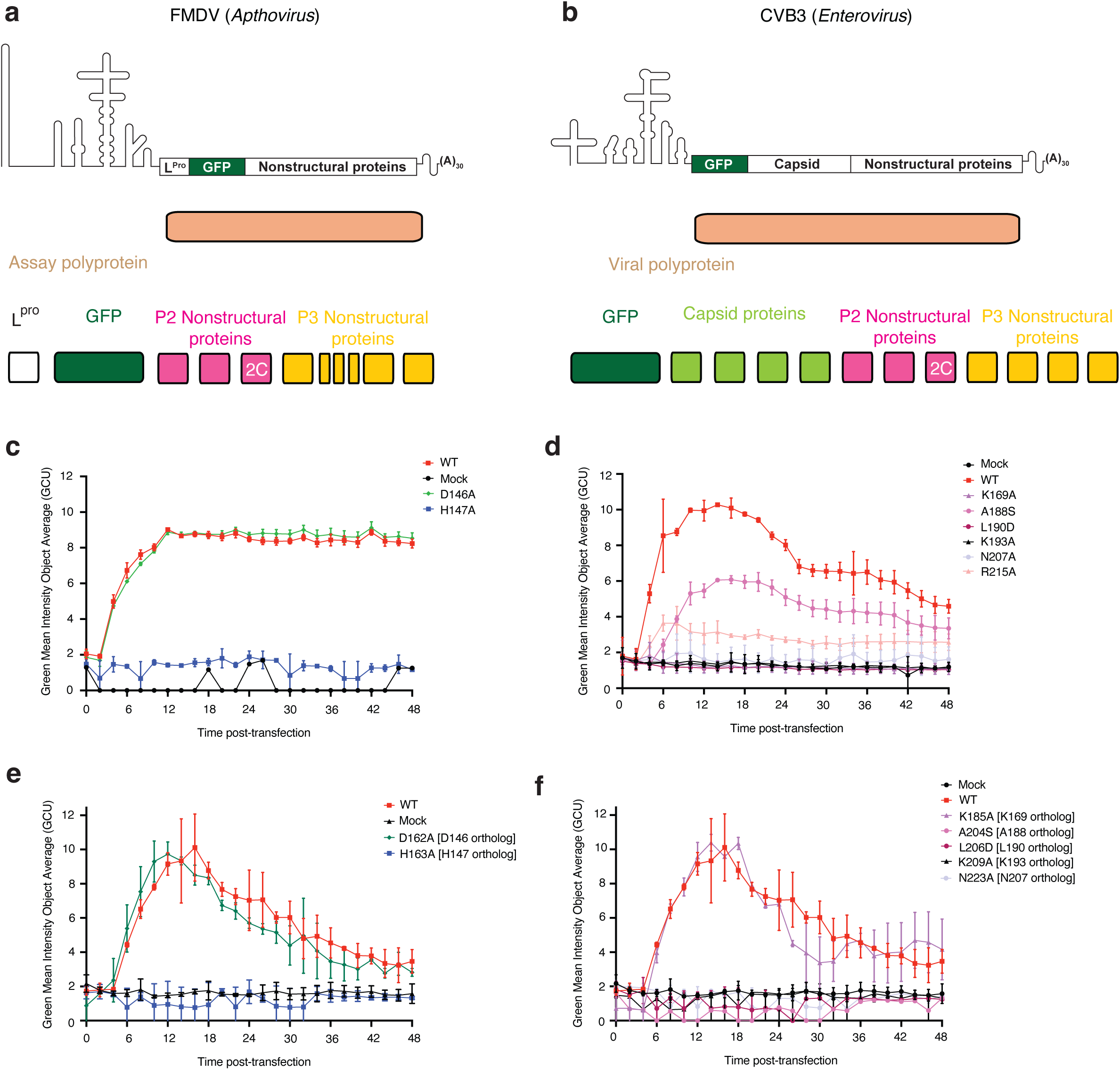
FMDV and CVB3 replication assays reveal functional residues of 2C: **a.** Schematic of FMDV replicon. Structural proteins are replaced by GFP. **b.** Schematic of CVB3 replicon. Viral protease cleavable GFP inserted upstream of structural proteins. **c-f.** Live cell imaging of GFP accumulation over 48-h time course following transfection of indicated RNAs. Mock-cells transfected with no RNA. **c and d.** FMDV WT and 2C mutant replicon GFP accumulation. Data in c and d were collected in different runs and corresponding WT are included in each. **e and f.** CVB3 WT and 2C mutant virus GFP accumulation. CVB3 2C D162A and H163A are shown separately in e for clarity, WT and Mock data are the same in e and f. c-f. Data are average of technical duplicates (error bars are SD) and representative of 3 biological replicates. Repeats are shown in **Extended Figure 9**.

### 2C is reminiscent of a AAA+ protein translocase rather than an SF3 helicase

2C has historically been classified as a clade 4 SF3 helicase based on sequence analysis^46^. Indeed, all clade 4 AAA+ proteins are of viral origin, just as 2C is^19^. However, given the conformation of our structure of the 2C:RNA holoenzyme complex, and its striking similarity to protein translocases, we performed a structural search against other AAA+ solved or predicted monomers (**Figure 5a**). We found that 2C clustered both near AAA+ SF3 helicases and protein translocases. Comparing the cryoEM solved complexes representative of these two hits, a member of the Hsp100 protein translocase family^47^ and SV40 T-antigen SF3 helicase^48^, we found that the overall arrangement of the 2C more closely resembled that of Hsp100 AAA+ protein translocases and other AAA+ protein translocases, forming a hexagonal arrangement around the substrate while SV40 and other SF3 helicases arrange their protomers into a star like arrangement^48–50^ (**Figure 5b**).

**Figure 5:**
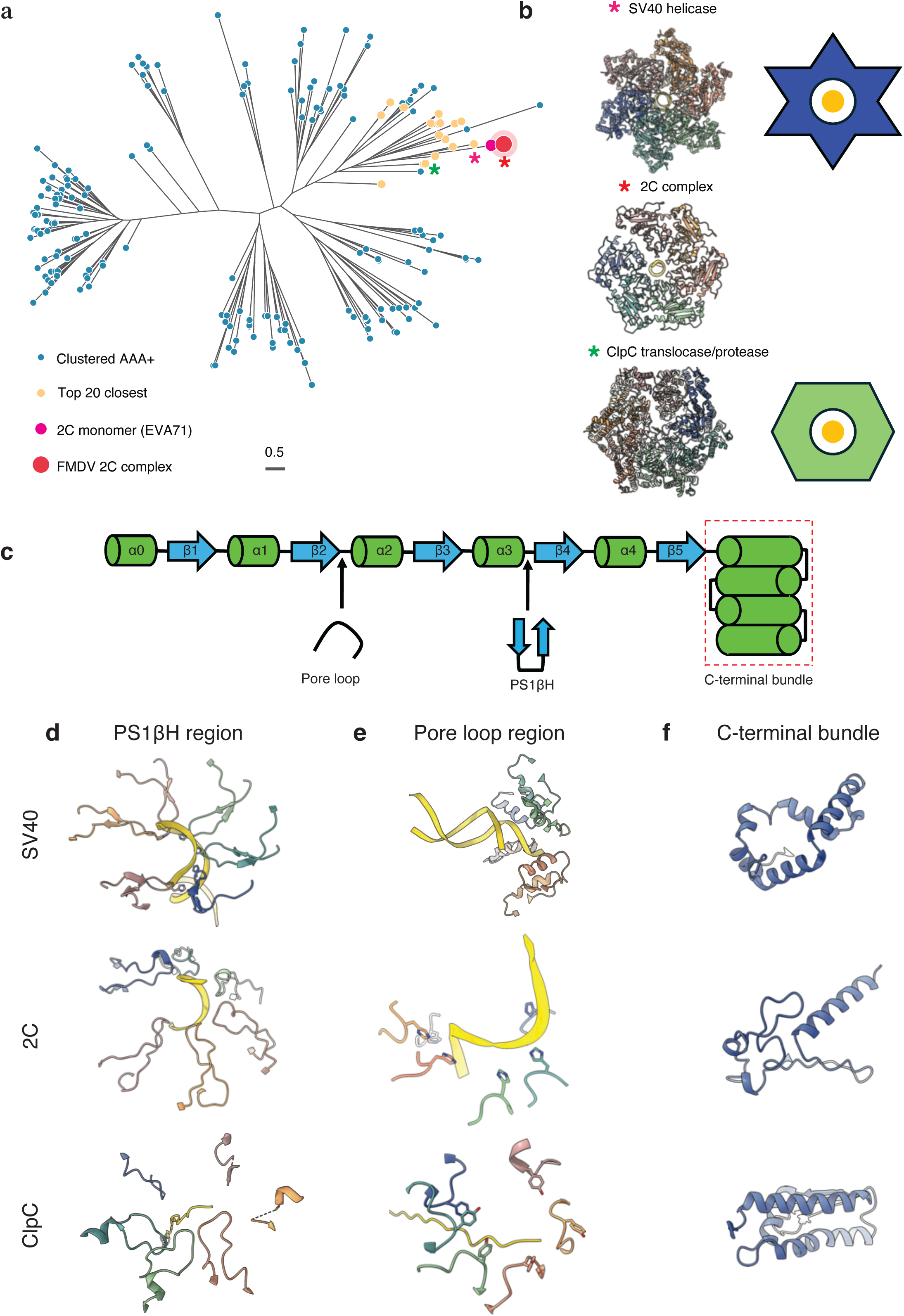
Structural phylogenetics of 2C relative to the AAA+ Superfamily: **a.** Unrooted phylogenetic tree of results from AAA+ structural search with the top 20 closest hits colored yellow, the EVA71 2C crystal structure colored pink and the FMDV 2C complex solved here as red. Hsp100 translocase (green star), SV40 SF3 DNA helicase (pink star) and FMDV-2C hexamer (red star) are indicated. **b.** Comparison of the hexameric structures of SV40 SF3 DNA helicase (PDB: 9KAE), 2C-RNA complex (this study), and a Hsp100 translocase (PDB: 6W24). Colored stars are as in a. The blue star with a golden circle (substrate) within it represents the overall conformational arrangement of typical AAA+ helicases, while the green hexagon with a golden circle (substrate) represents the typical arrangement of a protein translocase **c.** Linear schematic of a AAA+ domain. Pore loop insertion and PS1βH adaptations are shown as well, with a red box around the variable C-terminal bundle. Comparison of SV40, 2C, and Hsp100 translocase structures at the **d**. PS1βH region, **e.** pore loop region, and **f.** C-terminal bundle.

To more carefully examine 2C’s place within the AAA+ superfamily, we examined the key structural features that classify and define these proteins (**Figure 5c**). Specifically, AAA+ protein translocases typically engage substrate at a pore loop with a bulky aromatic amino acid that occurs between β2 and α2 while SF3 helicases are defined by a well ordered β-hairpin (PS1βH) that occurs between α3 and β4 that contains an aromatic amino acid that engages the DNA substrate^19^. Lastly, the C-terminal bundle region in SF3 helicases is uniquely rearranged compared to other AAA+ complexes and serves as a distinguishing feature. Looking first at the PS1βH, we unexpectedly find that SV40 T-antigen contains a well ordered β-hairpin with a histidine residue. Conversely, Hsp100 translocase and 2C do not contain a well ordered β-hairpin at the same position within the AAA+ topology and do not contain an aromatic residue (**Figure 5d**). At the pore loop region, SV40 T-antigen does not have any loop like feature nor aromatic residues whereas 2C and Hsp100 translocase both contain flexible loops with an aromatic amino acid intercalating the substrate’s backbone (**Figure 5e**). Lastly, 2C’s C-terminal region is relatively small and featureless as compared to both SV40 T-antigen and Hsp100 translocase (**Figure 5f**).

## Discussion

Here we present the structure of the picornaviral 2C:RNA holoenzyme complex revealing the complex arrangement of a longstanding target for the development of antiviral therapies. Like other AAA+ superfamily clade 4 SF3 viral helicases that unwind DNA, 2C binds to ssRNA in its central pore (**Figure 1c**).

We find the complex’s protomers in a variety of different nucleotide states (**Figure 1f-h**) yet all its protomers are nearly identical to each other (**Extended Figure 4f**). This is suggestive that the protomers translocate RNA substrate not through loop motions but with rigid body motions of the protomers, with a P_i_ release coupled to weakening of the interprotomer interface and ultimately with protomer translocation across the split^26^. The ADP nucleotide state of the protomers exists at the bottom and the top of the spiral. In AAA+ protein translocases, the bottom protomer hydrolyzes ATP into ADP and then rebinds at the top of the spiral staircase and exchanges nucleotide i.e. protomer F becomes protomer A and so on. However, biochemical evidence of 2C’s ability to translocate RNA is sparse. While one article reports helicase activity of 2C activity^2^, no other studies have been able to recapitulate helicase or translocase activity in any other 2C ortholog or biochemical condition^38,51–53^. Perhaps 2C, which assembles at the replication organelle, requires additional and presently unknown viral and/or cellular cofactors to be fully biochemically active. However, using our complex structure, we were able to separate RNA binding from function by comparing our RNA binding assays to viral replication assays. Indeed, the only mutant that did not show reduced ssRNA binding but completely ablated viral replication and is phylogenetically conserved was H147 (**Figure 3b**, **Figure 4c,e, Extended Figure 8b, Extended Figure 9**). Furthermore, H147 is located at the pore loop region where other AAA+ translocases in a variety of other systems contain an aromatic amino acid that engages the back bone of substrate to promote translocation of substrate. These results provide indirect but suggestive evidence that 2C likely translocates RNA to some degree. Furthermore, we found that the full length 2C only moderately increased binding to ssRNA (**Extended Figure 10**), indicating that the AAA+ domain is what primarily binds to substrate.

In this work we also characterize the specific modality employed by this AAA+ protein to bind RNA. A triad of amino acids, A188, L190, and K193 are the most biochemically critical and phylogenetic conserved residues that contribute to RNA binding (**Figure 2a-b**, **Figure 3d-f, h-j**, **Figure 4d,f, Extended Figure 6f,g, Extended Figure 8d-e, Extended Figure 9**). While mutations to K169 and R215 do ablate RNA binding in-vitro and lead to reduced viral replication in the FMDV replicon system, the lack of conservation of K169 and R215 and lack of effect of K169 in the CVB3 viral replication system (R215 was not conserved in CVB3) suggest that their relevance is more narrowly restricted to a smaller set of viruses within picornaviruses.

Previous work has suggested that 2C employs a two-step mechanism for binding^38^. The first step mediates binding of nucleic acid to 2C; 2C can bind RNA, DNA and modified nucleic acid polymers. In the second step, 2C proofreads the nucleic acid bound to ensure that it is RNA as RNA is the only polymer that can stimulate its ATPase activity. Our work here provides a structural basis for this two-step binding. L190 and K193, and additionally in FMDV K169 and R215, bind either to the phosphate backbone or base portion of the ssRNA. However, it is A188 carbonyl oxygen that confers 2C’s proofreading ability as it is the only specific interaction with the 2′-OH that is present in the 2C:RNA holoenzyme. DNA would be unable to make this interaction as it lacks a 2′-OH. This is reminiscent of the selectivity filter within the tetrameric pore of ion channels but instead the carbonyl oxygens line the inside of a hexamer in a staircase fashion rather than the pore of the channel^54^.

2C has been proposed to be multifunctional with one putative function of 2C being packaging of RNA within the capsid^15–17^. It is possible that 2C employs RNA translocation to encapsidate the viral RNA genome into the capsid to package the RNA. This would be analogous to AAA+ protein translocases that move protein substrate into different compartments^55,56^. Additionally, we show that 2C and its key residues responsible for binding and potentially translocating RNA are not limited to picornaviruses. Indeed we find a variety of undercharacterized 2C like orthologs in many other non-picornavirus RNA-virus families (**Extended Figure 7**). Notably, we show that norovirus’ NS3 contains many of the same binding and functional moieties that 2C does (**Figure 3h-j**).

The characterization of this 2C:RNA holoenzyme will accelerate drug development against 2C. Many putative inhibitors have been proposed and studied for 2C but they have relied on the monomeric crystal structure of a single protomer^3–5^. which lacks the experimental details of oligomerization that the holoenzyme presents. Our findings offer expanded potential for small molecule design and/or peptide inhibition that can underpin development of broad-spectrum picornavirus inhibitors.

Finally, we show that 2C contains many structural features reminiscent of the AAA+ protein translocases, as opposed to the SF3 helicases. The only viral clade in the AAA+ superfamily is clade 4, the SF3 helicases. Our structural analysis shows that 2C clusters near both SF3 helicases and protein translocases. Perhaps 2C and other SF3 helicases shared a common evolutionary origin but 2C diverged before other SF3 helicases acquired their helicase specific features. Alternatively, over the course of viral evolution, 2C potentially lost its SF3 helicase features and gained/adapted translocase-like features to support its function. Either way, our data demonstrates that 2C may be considered to occupy its own, novel AAA+ clade as an RNA-binding AAA+ protein.

---

**Extended Figure 1:**
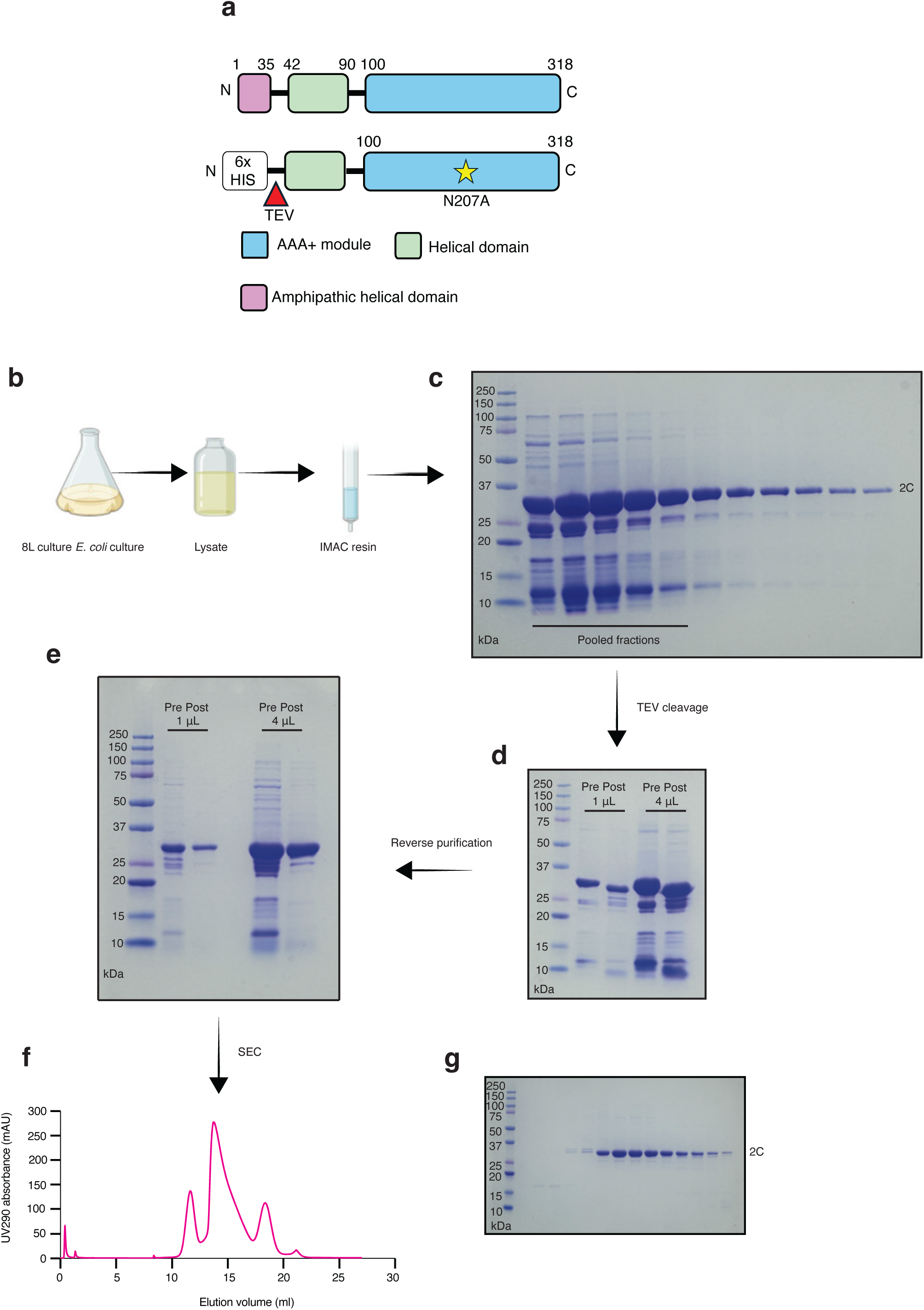
Purification scheme of 6x-HIS TEV Δ33 2C N207A: **a.** Schematic of FMDV 2C after polyprotein processing (top) and construct layout (bottom). **b-g**. Workflow of purification of Δ33 2C N207A. **b.** 8L cultures were induced with IPTG, lysed and subjected to affinity chromatography. **c.** SDS-PAGE gel of elution fractions. **d.** SDS-PAGE gel of protein before and after TEV cleavage treatment. **e.** SDS-PAGE gel of protein after subject to reverse affinity chromatography to remove cleaved 6x-His tag and other contaminants. **f.** SEC profile after application of protein. **g.** Final elution fractions for concentration after SEC.

**Extended Figure 2:**
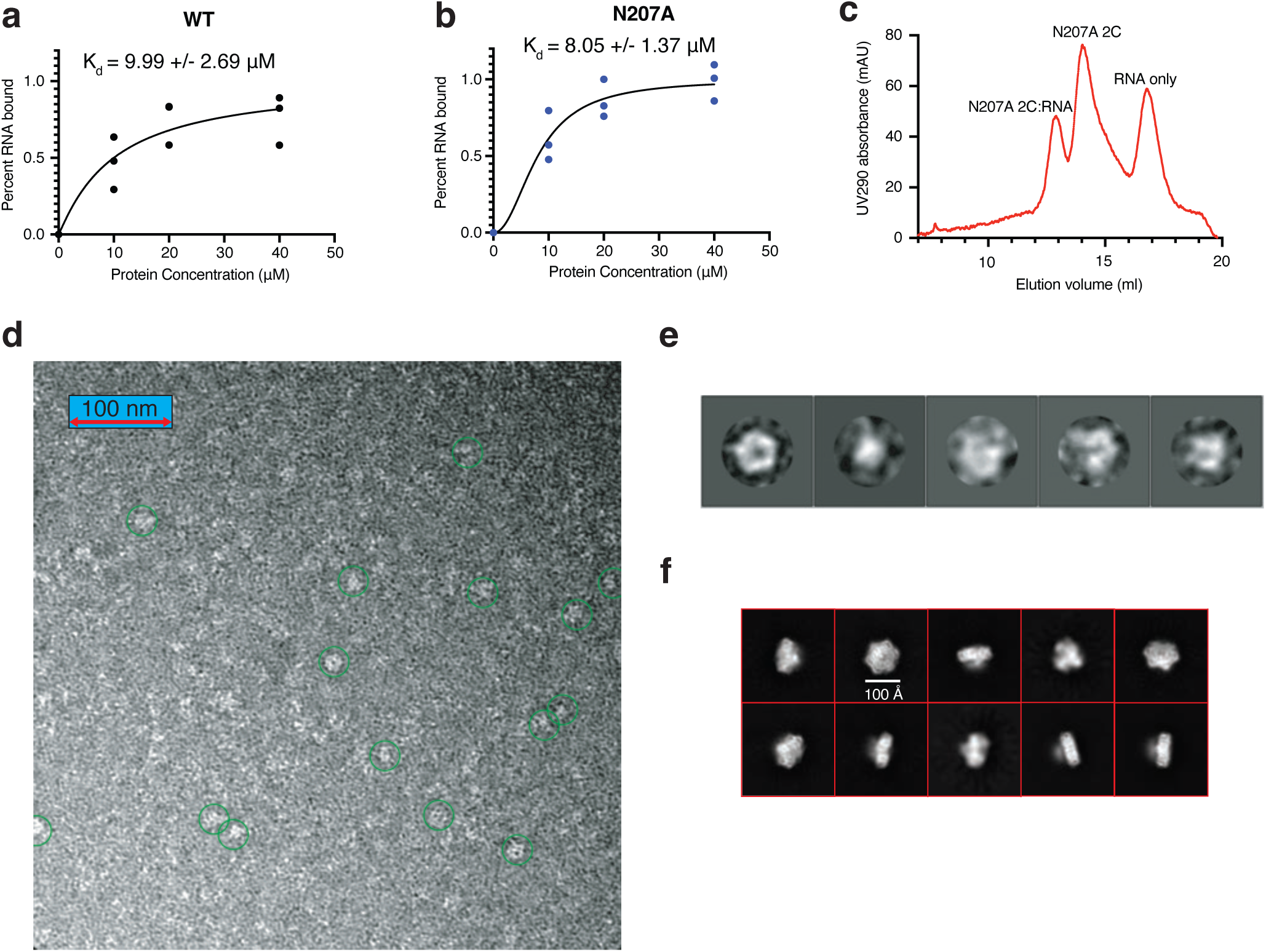
Biochemical interrogation and initial structural characterization of 2C: **a.** EMSA binding data (n=3) for Δ33 2C WT to ssRNA. **b.** EMSA binding data (n=3) for Δ33 2C N207A to ssRNA. **c.** SEC profile of 2C:RNA mixture after injection onto SEC. **d.** Representative image from negative stain electron microscopy with putative complexes in green circles. **e.** 2D class average of picked particles from representative micrographs from negative stain electron microscopy. **f.** 2D class averages from final particles used to reconstruct the final 3D volume of the 2C:RNA holoenzyme complex.

**Extended Figure 3:**
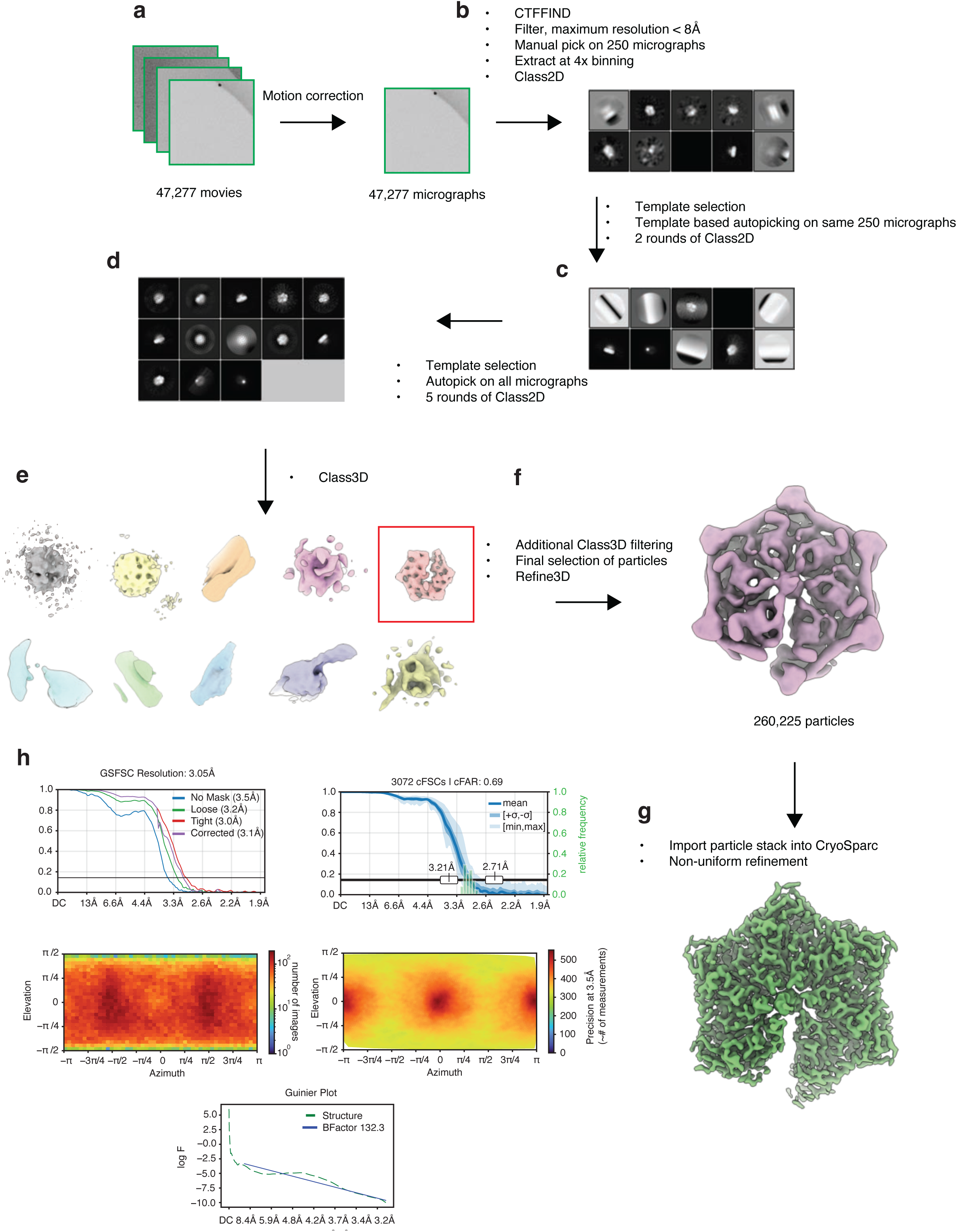
Processing schema of cryoEM data: **a.** 47,277 movies were initially collected and subject to standard preprocessing in Relion. **b-d.** Three cycles of picking, extraction and class2D were executed on larger pools of the micrographs until all micrographs were picked on. **e.** Initial class3D volumes with the red box representing class subjected to further classification. **f.** Low resolution volume representing final set of particles. **g.** Final sharpened map after additional processing in CryoSparc. **h.** cryoEM map statistics and reporting information.

**Extended Figure 4:**
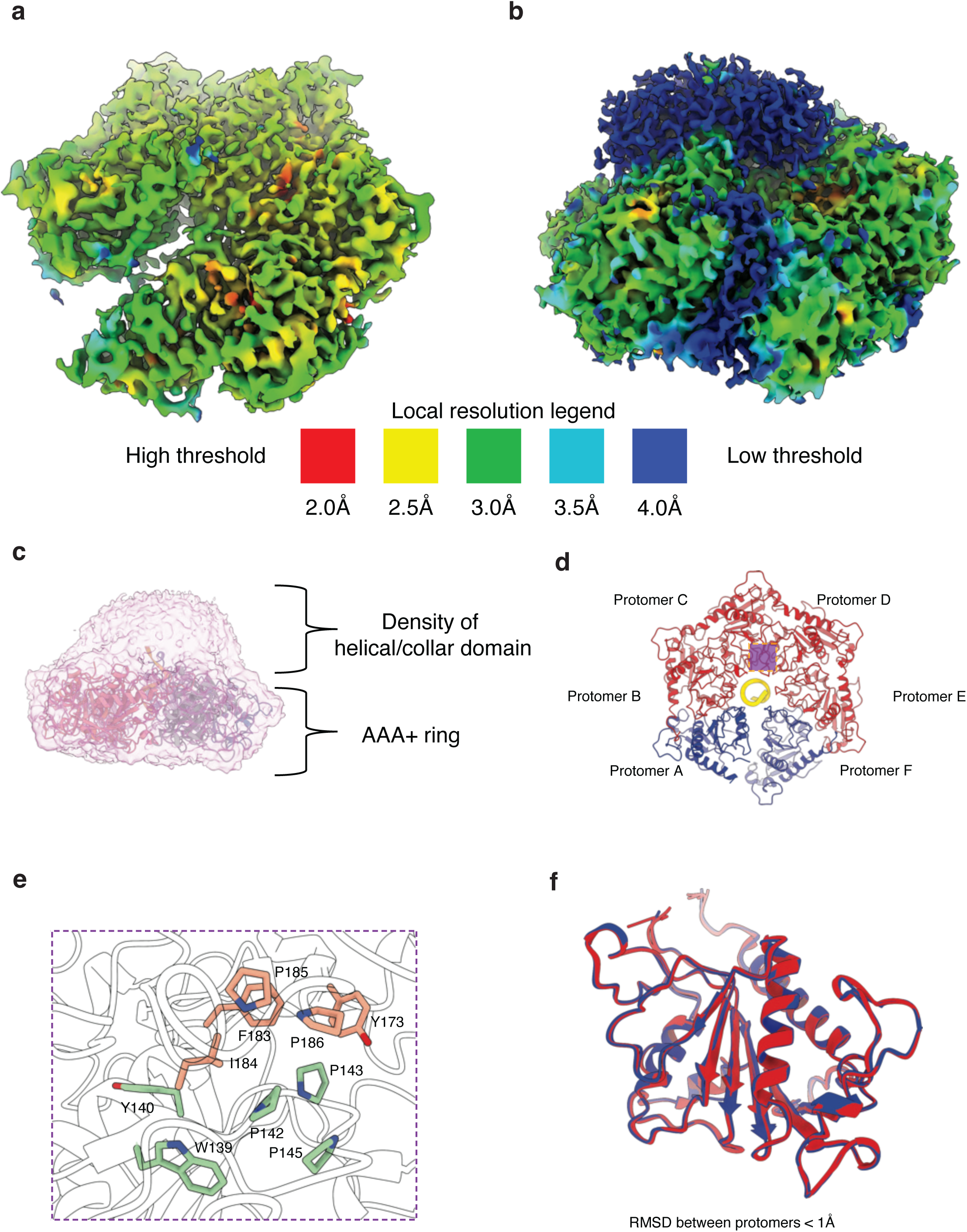
Local resolution and additional structural analysis of 2C: **a.** High threshold cryoEM map colored by local resolution. **b.** Low threshold cryoEM map colored by local resolution. **c.** cryoEM density of the presumably flexible collar/helical domain. **d.** Model of 2C:RNA holoenzyme colored by nucleotide state (blue: ADP, red:ATP). **e.** Purple shading represents hydrophobic depicted in **e.** Hydrophobic pocket at the interface of protomers is depicted, shaded purple in d. **f.** Alignment of protomers in different nucleotide states.

**Extended Figure 5:**
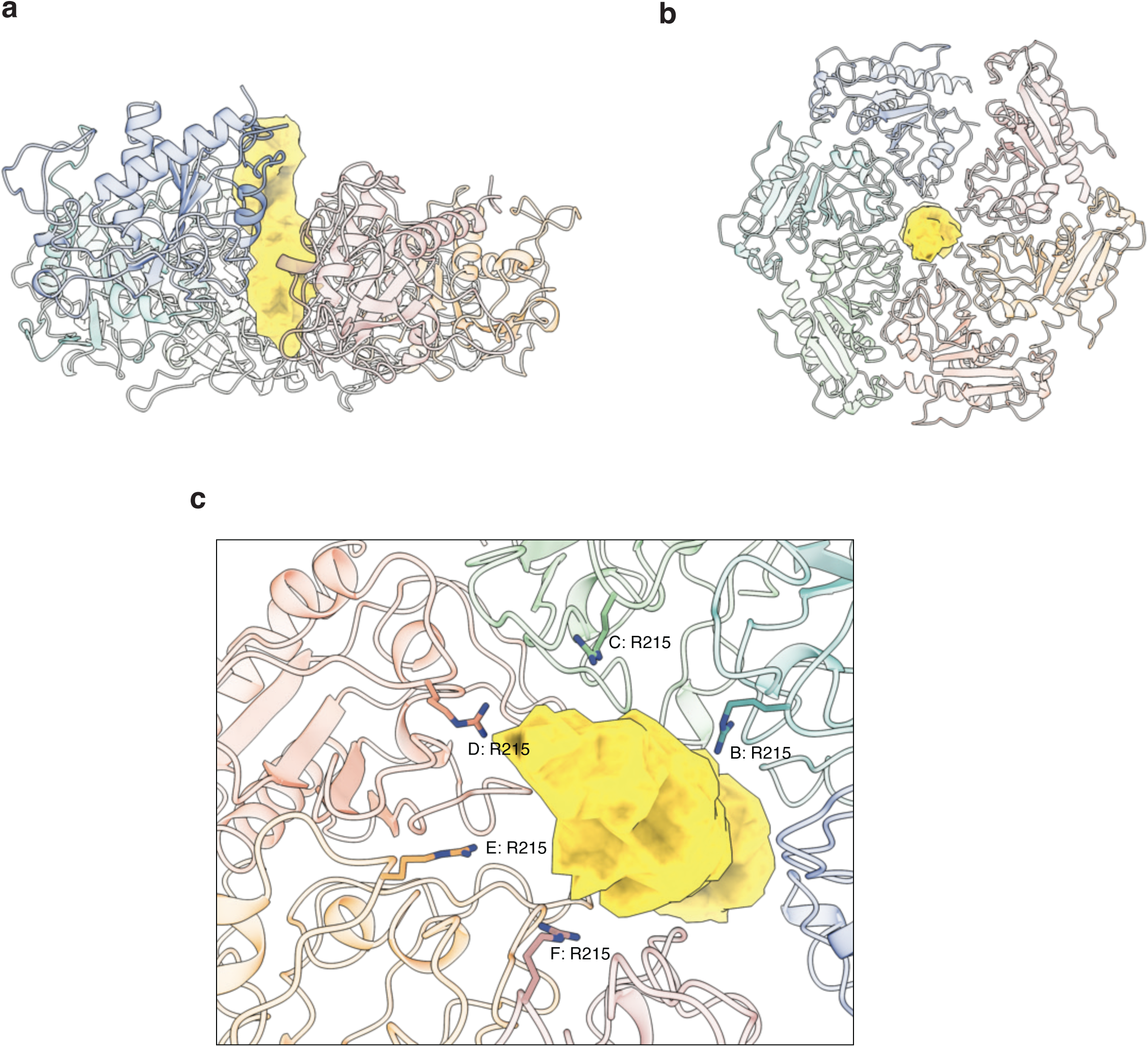
Extended ssRNA density in the pore of 2C: **a.** Side view of unsharpened cryoEM density of density corresponding to extended ssRNA (gold) in the pore. **b.** Top view of the same density in **a**. **c.** Zoomed view near the bottom of the 2C:RNA holoenzyme complex depicting R215 from each protomer within hydrogen bonding distance of the ssRNA density.

**Extended Figure 6:**
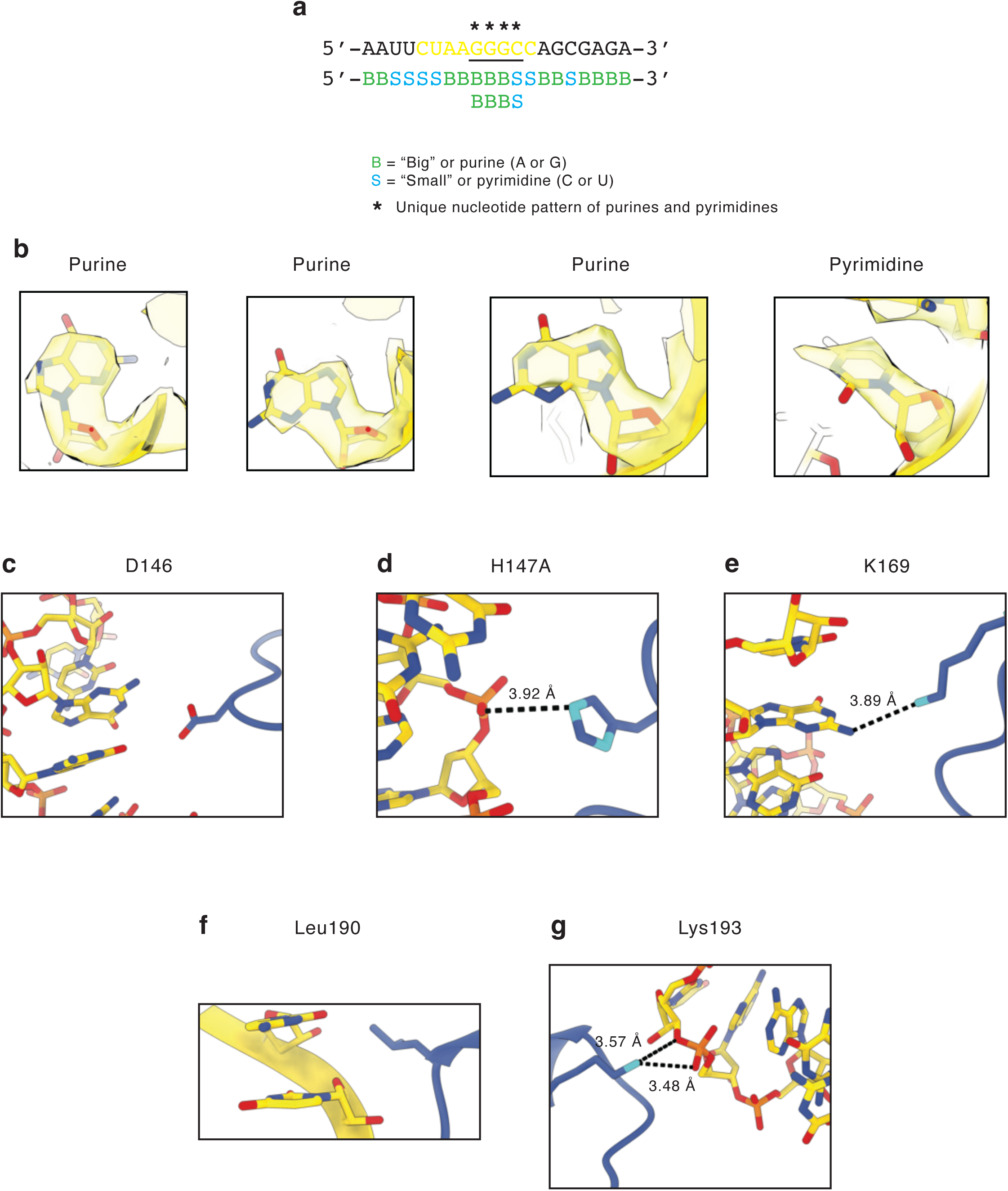
Indexing of ssRNA and examination of pore residues: **a.** Sequence of the ssRNA used in cryoEM labeled as “big” or “small” depending on whether it is a purine or pyrimidine, respecitvely. Nucleotides with a * above them indicate the unique motif used to register the nucleotides. **b.** CryoEM density of GGGC region of ssRNA. **c.** D146 has no proposed direct interactions with ssRNA. **d.** H147 can potentially form a weak hydrogen bond to the phosphate backbone. **e.** K169 can potentially form a weak hydrogen bond with the Watson-Crick face of the nucleotide. **f.** Leu190 packs against the hydrophobic base stacking face. **g.** K193 can hydrogen bond to multiple locations on the ssRNA backbone.

**Extended Figure 7:**
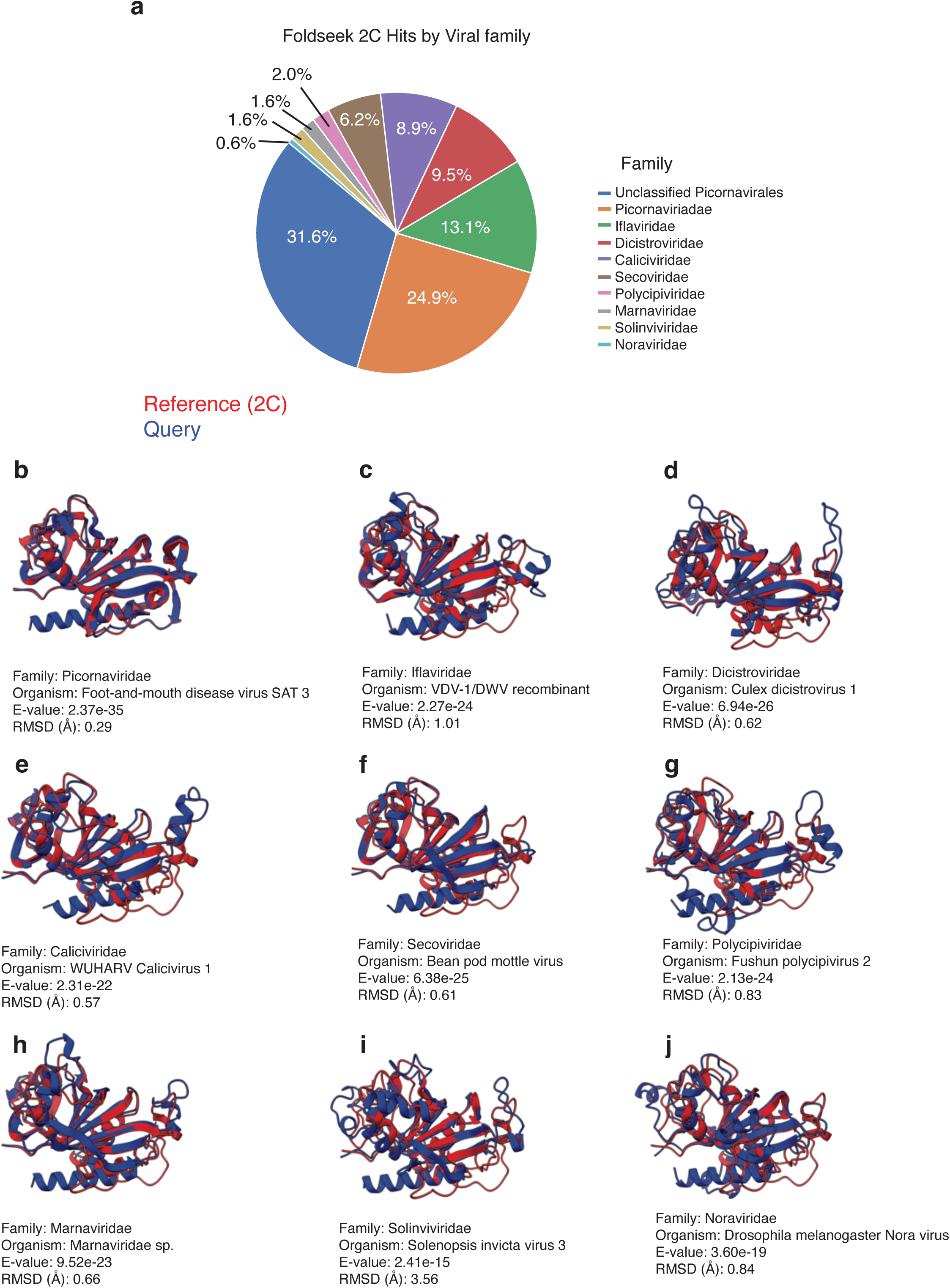
2C orthologs in Picornavirales: **a.** Pie chart of identified hits of FoldSeek search against Picornavirales showing the percentage of hits (above E-value of 0.001) that bin into different families. **b-j.** Comparison of top hit per family (as assessed by E-value) where structures colored red represent the query structures and the blue structures represent the predicted structures from **b.** *Picornaviridae*, **c.** *Iflaviridae*, **d.** *Dicistroviridae*, **e.** *Calciviridae*, **f.** *<u>Secoviridae,</u>* **g.** *Polycipiviridae*, **h.** *Marnaviridae*, **i.** *Solinvivirdae*, **j.** *Noraviridae*.

**Extended Figure 8:**
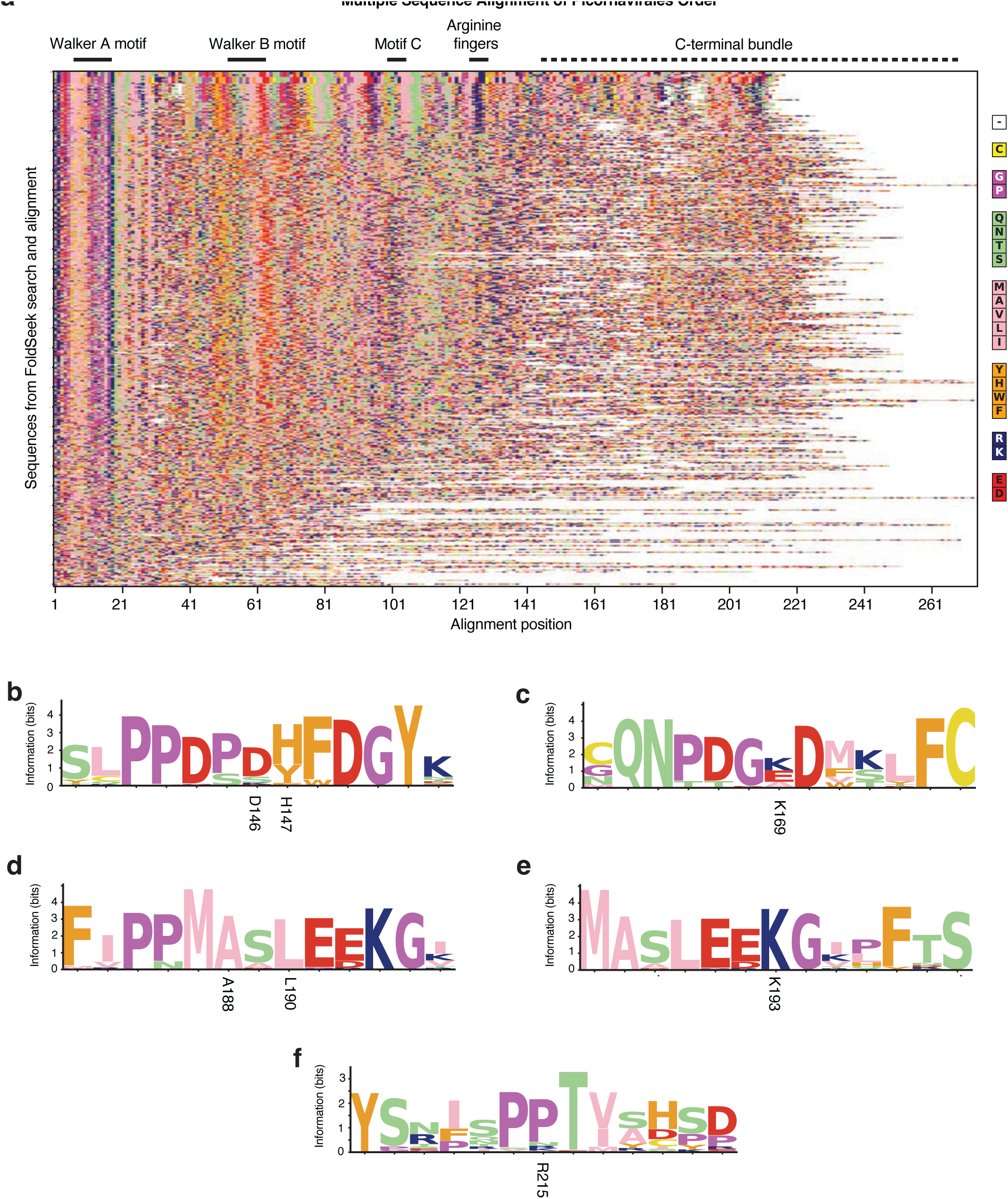
Alignment of deep Picornavirales orthologs: **a.** Heatmap alignment of hits from Foldseek queried across Picornavirales. Each row represents a unique sequence:structure pair and the top sequence:structure pair is the FMDV 2C structure. The alignment is based on the structure and thus begins at the beginning of the model. The legend on the right reflects the coloring scheme for the amino acids sorted by chemical property. For b-f, a representative set of sequences were taken to generate interepretable sequence logos. **b.** Sequence logo alignment zoom in at D146 and H147. **c.** Sequence logo alignment zoom in at K169. **d.** Sequence logo alignment zoom in at A188 and L190 region. **e.** Sequence logo alignment zoom in at K193. **f.** Sequence logo alignment zoom in at R215.

**Extended Figure 9:**
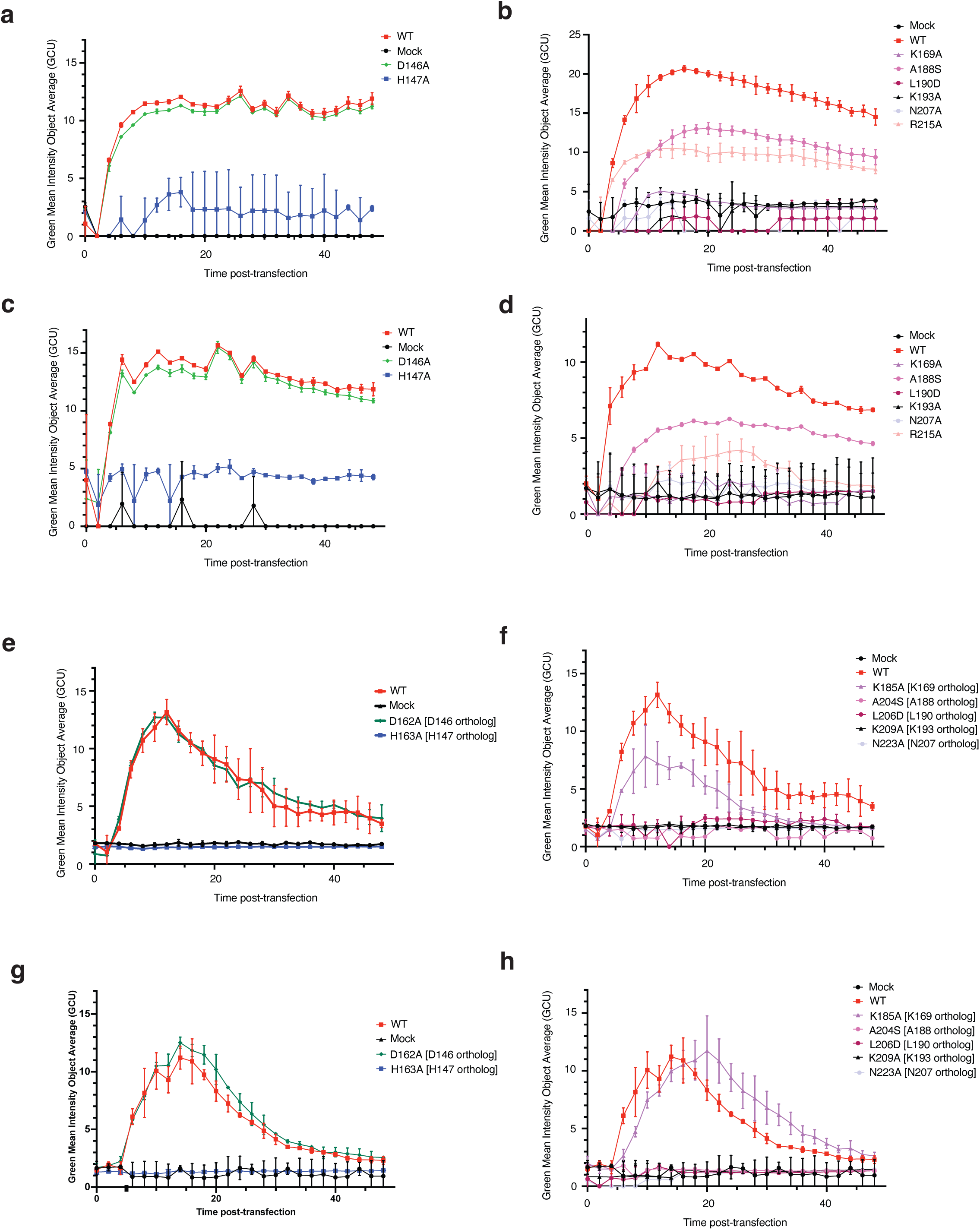
Additional replicon assays of 2C: **a-d.** Biological replicates of FMDV replicon data shown in main Figure 4 c-d. **e-h.** Biological replicates of CVB3 virus data shown in main Figure 4 e-f. Data are averages of two technical replicates (error bars are SD).

**Extended Figure 10:**
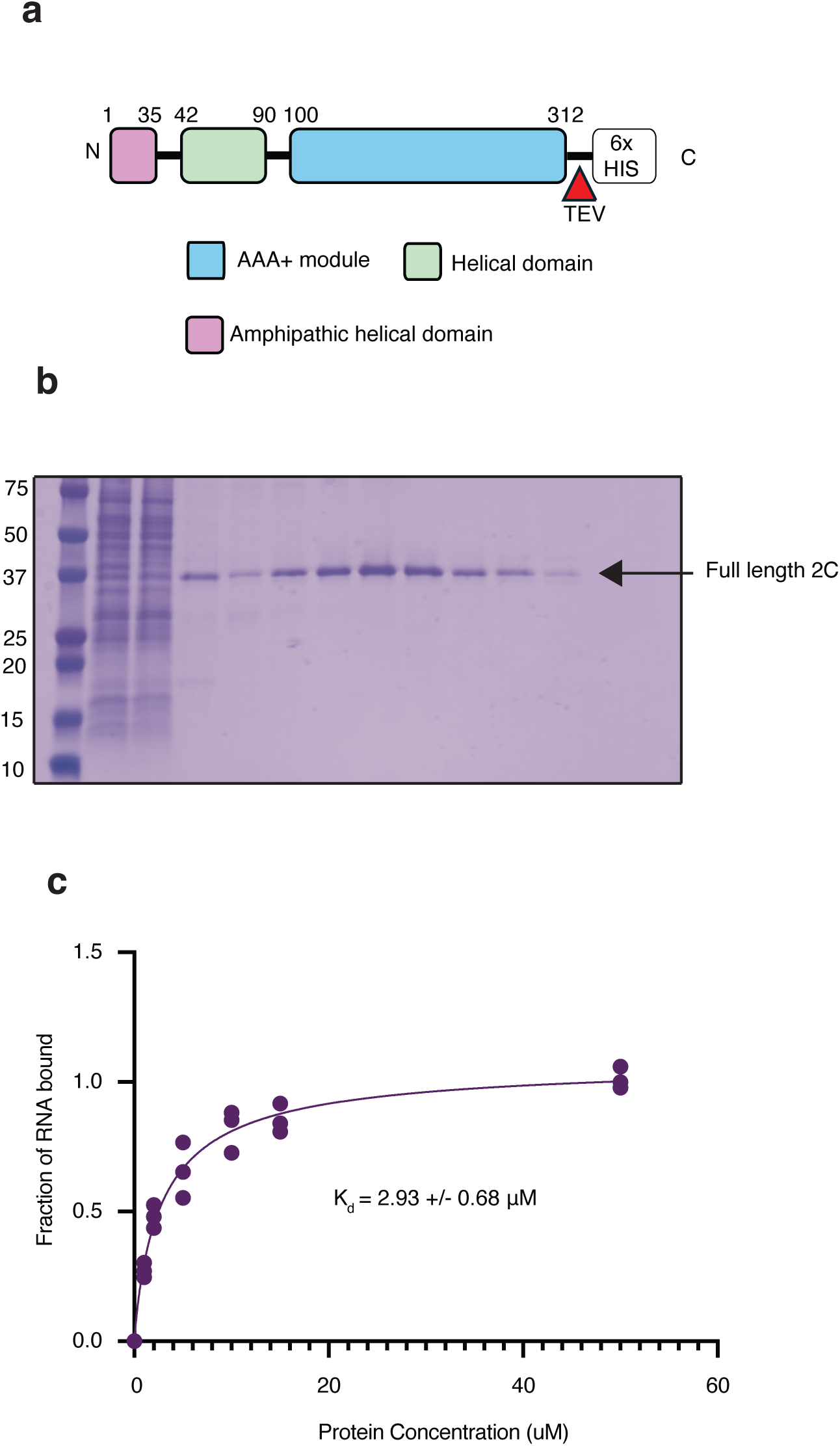
Full length FMDV 2C: **a.** Schematic of construct used to purify full length FMDV 2C. **b.** Final gel of full length 2C after SEC. **c.** EMSA experiments to evaluate binding of FL FMDV 2C to ssRNA.

## Materials and Methods

### Preparation of the 2C:RNA holoenzyme in hydrolyzing conditions

A construct containing 6x-HIS TEV Δ33 2C N207A^23^ was transformed into BL21 DE3 cells. 8L cultures were grown and cells were induced to express protein. Cell pellets were resuspended in a lysis buffer of 50 mM HEPES pH 7.1, 400 mM NaCl, 1 mM β-mercaptoethanol, and 0.1% Triton X-100. Following lysis, lysate was clarified with a 40,000g spin. The supernatant was then bound to IMAC resin, washed with lysis then eluted with imidazole. Fractions were collected and cleaved by TEV protease. The protein was then subjected to reverse affinity-chromatography and then subjected to size exclusion chromatography (SEC) with a SEC buffer of 50 mM HEPES pH 7.1, 200 mM NaCl, 1 mM β-mercaptoethanol. ssRNA (5′-AAUUCUAAGGGCCAGCGAGA-3′) was purchased directly from IDT DNA. 2C protein and ssRNA were mixed and subjected to SEC in the final protein buffer supplemented with MgCl_2_ and ATP. The peak corresponding to the 2C:RNA protein holoenzyme was used for further structural studies.

### cryoEM of the 2C:RNA holoenzyme in hydrolyzing conditions

Quantifoil R1.2/1.3 200-mesh gold grids were treated with chloroform and dried overnight. Grids were glow-discharged and 4 μl of the sample with 0.05% v/v Nonidet P-40 was blotted onto the grids and further blotted and vitrified in liquid ethane using an FEI Vitrobot (Thermo Fisher Scientific). Grids were blotted for 4 s with a 5-s wait time. The single-particle cryoEM data collection and processing workflow are described thoroughly in **Extended Figure 3.** An FEI Titan Krios (Thermo Fisher Scientific) cryo-EM instrument with a Falcon F4i camera was used for data collection. Micrographs were analyzed using a combined workflow of Relion 5^57^ and Cryosparc 4^58^. Models were built using a combination of AlphaFold3^59^ as an initial prediction for subdomains of a single protomer with further refinement and complex building in COOT^60^ and phenix^61^. ChimeraX was used for visualization and analysis of data.

### Electrophoretic gel shift assays (EMSAs)

Increasing concentrations of WT or mutant 2C protein were mixed with fluorescent 100 nM ssRNA labeled with 5′ 6-FAM in a final binding buffer of 50 mM HEPES pH 7.1, 40 mM NaCl, 2 mM MgCl_2_, and 0.5 mM TCEP. Protein and RNA mixtured were loaded shortly after mixing onto Any kD Mini-Protean gels (Bio-Rad, 4569033) and separated under native conditions with Tris–glycine running buffer (Bio-Rad), since varying incubation times had no difference on the binding. Gels were directly imaged on an iBright 1500.

### FMDV replicon and CVB3 virus

Plasmids containing the FMDV-GFP replicon and CVB3-GFP reporter virus were modified by insertion of fragments containing desired 2C mutations generated by Twist Bioscience and confirmed by sequencing. Plasmids were linearized and used as templates for *in vitro* transcription of RNA. Baby hamster kidney (BHK-21) cells were seeded at a density of 1.5 × 10⁴ cells per well in 96-well plates one day prior to transfection. Transfections were performed using 0.5 µg of *in vitro*-transcribed RNA diluted in jetMESSENGER® mRNA transfection buffer and 1µL of jetMESSENGER® (Sartorius, 101000005). Images of transfected cells were captured every 2 h for 48 h using an Incucyte S3 Live Cell Imaging system (Sartorius) housed within an incubator maintained at 37 °C and 5% CO₂. Images were captured from one or two regions/well using the 10× objective. GFP fluorescence mean intensity was quantified using IncuCyte 2025C image processing software and used as a surrogate measure of viral RNA replication kinetics. All experiments were performed using three independent biological replicates, each including technical duplicates.

### Structural phylogenetic analysis of AAA+ ATPase Domains

PDB structures annotated with the AAA+ ATPase (IPR003593) or SF3 helicase (IPR014015) domains were retrieved from InterPro^62^ (accessed May 2026) via its REST API, yielding 15,279 domain annotations across 3,431 unique entries. Structures were downloaded from the RCSB PDB in mmCIF format, and for each entry the annotated domain was extracted, retaining the single chain with the most standard amino acid residues for oligomeric assemblies; this produced 3,431 single-chain domain sequences. Redundancy was reduced with MMseqs2^63^ easy-cluster (v18.8cc5c; 40% identity, 80% bidirectional coverage, --cov-mode 0 --cluster-mode 1) to 197 non-redundant representatives, which together with the query structure (2C chain C) were aligned using FoldMason^64^ easy-msa. FoldMason’s progressive alignment over both the 3Di structural alphabet and amino acid sequences is robust to the low sequence identity (<20%) typical of remote AAA+ homologues, yielding an alignment of 198 sequences across 1,947 columns. Maximum likelihood phylogenetic inference was performed with FastTree^65^ (v2.1.11) under the JTT substitution model with a 20-category gamma distribution (+Γ), and branch support was assessed using SH-like local supports. Trees were rendered as unrooted radial phylograms (Biopython v1.87, matplotlib v3.10) with branch lengths proportional to expected substitutions per site; a small number of extreme outliers (>95th percentile of branch lengths) were capped for display only, without affecting the reported topology or branch lengths.

### Structural phylogenetic search of 2C orthologs in Picornavirinae

The 2C structure was used as a query for FoldSeek^66^ server against the BFVD^67^ in 3Di/AA with a taxonomic filter set to *Picornavirales* which yielded 548 hits, 497 of which were below the default e-value threshold of 0.001. Following this, the taxonomic information of the significant hits were extracted to generate statistics on the distribution of 2C orthologs in other *Picornavirales* families. The best hit of each family was taken and the generated c-α alignment of the core region was used to generate representative structural alignments between the query and found structures.

### Purification of full length FMDV 2C

The full length sequence of 2C tagged was tagged with 6x-HIS tag prior to it being cloned in pFastBac and expressed in SF9 cells using the Bac-to-Bac expression system (ThermoFisher). FMDV 2C full-length protein was expressed in Sf9 insect cells and purified from a 2L frozen cell pellet in 50 mM HEPES pH 7.5, 400 mM NaCl, 0.5 mM TCEP. Membranes were solubilized in 2% DDM at 4°C, clarified by ultracentrifugation, and the supernatant was batch-bound to resin overnight. Protein was eluted with 300 mM imidazole and further resolved by SEC in 20 mM HEPES pH 7.5, 100 mM NaCl, 0.02% DDM, 0.5 mM TCEP. Following TEV digestion and reverse chromatography to remove the His-tag, the purified protein was dialyzed overnight and concentrated in a 100K MWCO spin concentrator.

### Negative stain transmission electron microscopy

Grids were prepared for negative stain transmission electron microscopy using Formvar/carbon-coated grids. Grids were glow-discharged with the carbon side facing up immediately prior to use. A piece of parafilm was laid flat and secured with tape, and three droplets of ∼200 µL deionized water were arranged in a row per sample condition. Protein samples were prepared at two concentrations (0.025 mg/mL and 0.0125 mg/mL) and 3 µL of each was applied to the carbon side of a glow-discharged grid and incubated for 1 minute. Grids were then washed three times by floating carbon-side down on successive water droplets for approximately 30 seconds each. Grids were subsequently stained by floating on a droplet of uranyl acetate for 1 minute, blotted with Whatman paper, and allowed to air dry before storage. Negatively stained grids were imaged on a Morgagni transmission electron microscope operating at 100 kV. Images were acquired using a Gatan Rio9 camera in Gatan Digital Micrograph at a camera temperature of 10°C, with an applied defocus of −2.0 to −3.0 µm and exposure times up to 2 seconds.

